# Motile bacteria leverage bioconvection for eco-physiological benefits in a natural aquatic environment

**DOI:** 10.1101/2023.06.06.543831

**Authors:** Francesco Di Nezio, Samuele Roman, Antoine Buetti-Dinh, Oscar Sepúlveda Steiner, Damien Bouffard, Anupam Sengupta, Nicola Storelli

## Abstract

Bioconvection, the active self-sustaining transport phenomenon triggered by the accumulation of motile microbes under competing physico-chemical cues, has been long studied, with recent reports suggesting its role in driving ecologically-relevant fluid flows. Yet, how this collective behaviour impacts the ecophysiology of swimming microbes remains unexplored. Here, through physicochemical profiles and physiological characterizations analysis of the permanently stratified meromictic Lake Cadagno, we characterize the community structure of a dense layer of anaerobic phototrophic sulfur bacteria, and report that the associated physico-chemical conditions engender bioconvection when bulk of the motile purple sulfur bacterium *Chromatium okenii* synchronize their movement against the gravity direction. The combination of flow cytometry and fluorescent *in situ* hybridization (FISH) techniques uncover the eco-physiological effects resulting from bioconvection, and simultaneous measurements using dialysis bags and ^14^C radioisotope, allowed us to quantify *in situ* the diurnal and nocturnal CO_2_ fixation activity of the three co-existing species in the bacterial layer. The results provide a direct measure of the cellular fitness, with comparative transcriptomics data – of *C. okenii* populations present in regions of bioconvection vis-à-vis populations in bioconvection-free regions – indicating the transcripts potentially involved in the bioconvection process. This work provides direct evidence of the impact of bioconvection on *C. okenii* metabolism, and highlights the functional role of bioconvection in enhancing the metabolic advantage of *C. okenii* relative to other microbial species inhabiting the microbial layer.

## Introduction

Bioconvection is a well-known collective behavior observed in diverse groups of motile microorganisms, from bacteria, to algae, to protists, which share a common upward-swimming behavior and a density higher than water^1–3^. The convective motion is triggered when a large number of such microorganisms accumulates in a water body, leading to an increase in local density and the subsequent formation of a density-unstable fluid ambient. As a result, the denser mixture of water and microorganisms generates hydrodynamic instabilities with the underlying water whereby the denser cell-rich layer comes down due to the gravity force in the form of characteristic ‘plumes’. The bioconvection cycle is sustained by the microorganisms carried away from the sub-surface layer being replaced by others upswimming from below, generating a convective cycle^4–6^.

Microbial bioconvection in natural aquatic settings is triggered when up-swimming cells accumulate in competing gradients – physical or chemical – which get established due to the local processes^7^. While the focus of earlier studies on bioconvection has been on photo- and gyrotaxis^7, 8^, more recent works have indicated the possibility that microbes could harness bioconvection for eco-physiological advantage, particularly in quiescent aquatic environments, for instance in meromictic lakes. In general, however, the physiological and ecological implications of microbial bioconvection still remain to be clarified.

Meromictic lakes exhibit permanent density stratification and vertical gradients of light and redox conditions. Their water column hosts several physiological groups of microorganisms distributed along these vertical gradients, which act as specific ecological niches. These characteristics make them an exemplary field-scale laboratory for studying anaerobic microorganisms and their associated biogeochemical processes^9–11^. In the particular case of shallow chemocline (i.e. with depth scaling with the photic zone), light penetration can promote large microbial blooms throughout the chemocline, with for instance blue-green algae in its upper part and phototrophic sulfur bacteria in its lower part^12, 13^. Lake Cadagno is an iconic example of crenogenic meromixis, a phenomenon occurring when saline springs discharge dense water onto a freshwater lake depression, setting up density stabilizing conditions^14^. As a result, the water column is permanently stratified into an oxic, electrolyte-poor mixolimnion, a salt-rich, sulfidic monimolimnion and an intervening chemocline at 10.0 – 12.0 m depth with reverse oxidoreductive gradients, such as the disappearance of oxygen and the appearing of sulfide^15^.

One of the main features of Lake Cadagno is the diverse community of anoxygenic phototrophic sulfur bacteria that develops in the chemocline proximity characterized by opposite gradients of sulfide (S^2-^) and light irradiance^16–20^. This community forms a dense bacterial layer (BL) and is composed of purple sulfur bacteria (PSB) and green sulfur bacteria (GSB), a number of which have been successfully isolated and cultivated in the laboratory^21–24^. At the present day, PSB strains *Chromatium okenii* LaCa and *Thiodictyon synthophicum* Cad16^T^, and GSB *Chlorobium phaeobacteroides* 1VII D7, previously isolated from Lake Cadagno and grown as pure cultures in the laboratory, are the dominant phototrophic sulfur bacteria species in the BL. PSB and GSB play a major role in the biogeochemical cycles of carbon^25^, sulfur^26^ and nitrogen^27^. In particular, inorganic carbon fixation has also been observed in the absence of light, under microoxic conditions, especially in PSB^28, 29^.

Bioconvection can have important ramifications for microbes in natural settings, including molecular transport over large environmental scales, physico-chemical ecology and spatio-temporal distribution of microbes in aquatic ecosystems. Recently, by combining field and laboratory studies with numerical modeling, Sommer *et al*.^30^ demonstrated that the motile PSB *Chromatium okenii* is able to trigger mixing through bioconvection, resulting from the concerted movement of large portions of its population. Lake Cadagno was the first example of bioconvection witnessed in a natural freshwater environment^30^, so far limited to a few observations in marine ecosystems or laboratory settings^31–35^. However, the study by Sommer *et al*. focused on the physical demonstration of the phenomenon without exploring its consequences on microenvironmental conditions, microbial community, or biogeochemical implications on the lake ecosystem.

Regular monitoring of Lake Cadagno water column has shown the occurrence of bioconvection between June and late August^30, 36, 37^. In this study, we compare physico-chemical profiles and physiological characterizations at two different times of the summer season, July and September 2020, when we observed the presence and absence of bioconvection, respectively, to elucidate its eco-physiological effects on the BL microbial community. Additionally, we carried out transcriptomics analyses of PSB *C. okenii*, the actor of bioconvection, with the goal of identifying key genes involved in this process or, more generally, in its seasonal behavior. Our results imply that bioconvection may provide *C. okenii* with an ecological advantage over the coexisting phototrophic sulfur bacteria.

## Methods

### Study site and sampling

Lake Cadagno is located in the Piora Valley (46°33’N, 8°43’E) in the southern Swiss Alps. The sampling was conducted on 16 July and 17 September 2020 using a pump-CTD system (CTD115M, Sea & Sun Technology, Germany), as described in Di Nezio *et al*.^38^. For the physico-chemical characterization of the water column, *in situ* high vertical resolution data (sampling at 16 Hz) were obtained during a first continuous downcast of the CTD system from the lake surface down to ∼18.0 m depth. During a second downcast, after 30 min and in a different area, discrete water samples were collected from a total of 6 depths (between 12.0 m and 18.0 m) for bio-chemical analyses and from the top of the BL for incubation experiments (see ‘Radioisotope incubation and uptake analysis’ section).

Profiles were recorded at sunrise on 16 July and 17 September 2020 at 05:15 h and 06:30 h, respectively. CTD profiles for determination of light regimes were measured at daytime (17:00 h) on both days. Atmospheric radiation data at 10 min resolution for both sampling campaigns were retrieved from a meteorological station (istSOS; https://hydromet.supsi.ch/) close to the lake shore.

Water samples for microbiological (FISH and flow cytometry) and chemical (S^2−^, SO_4_^2−^ and CaCO_3_) analyses were stored in 50 ml falcon tubes and processed within the following hour, as described in Di Nezio *et al*.^38^.

### Cell growth

Phototrophic sulfur bacteria were grown in Pfennig’s medium I^39^ and cultivated in laboratory under a light/dark photoperiod of 16/8 h with a light intensity of 38.9 μmol m^-2^ s^-1^ PPFD (photosynthetic photon flux density) within the photosynthetically active radiation (PAR) range, measured with a portable LI-180 Spectrometer (LI-COR Biosciences, Lincoln, NE), as in Di Nezio *et al*.^38^.

### Bacterial layer microbial community analysis

To describe the composition of the BL microbial community, cell counting was performed through flow-cytometry (FCM), as described in Danza *et al*.^26^. Bacterial populations were distinguished by applying gates on cell size (forward scatter, FSC) of 0.1 - 1.0 μm for GSB, 2.0 - 4.0 μm for small-celled PSB and 4.0 - 10.0 μm for *C. okenii* (Fig. S1). Simultaneously, fluorescent *in situ* hybridization (FISH) analyses were carried out on bacterial layer (BL, zone with a turbidity > 10 FTU) water samples, as previously described in Decristophoris *et al*.^17^ (Tab. S1), spotting 2 μl of fixed sample on gelatin coated slides (0.1% gelatin, 0.01% KCr(SO_4_)_2_) and observing them by epifluorescence microscopy using filter sets F31 (AHF Analysentechnik, Tübingen, Germany; D360/40, 400DCLP, D460/50 for DAPI) and F41 (AHF Analysentechnik, HQ535/50, Q565LP, HQ610/75 for Cy3) at 100X magnification.

### Radioisotope incubation and uptake analysis

To test the photosynthetic efficiency of pure bacterial laboratory cultures, cells were grown up to concentrations of 10^6^ cells ml^-1^ (mid exponential phase), sealed in 50-cm-long dialysis bags (inflated diameter of 62.8 mm; Karl Roth GmbH Co. KG, Germany), whose membrane allows for diffusive transport of small molecules (< 20 kDa); thus, preventing contamination of the incubated samples from the surrounding environment.

The dialysis bags were acclimatized to the chemocline conditions of Lake Cadagno for a period of 4 weeks (from 15 June to 16 July 2020 and from 21 August to 17 September 2020) before the experiments, at varying depths between 12.20 – 12.40 m and 13.12 –12.94 m in July and September, respectively.

To measure the amount of light reaching the incubation depth, the dialysis bags supporting frame was equipped with HOBO UA-002–64 Pendant passive data loggers (Onset Computer Corporation, MA), one at the top and one at the bottom of the structure, measuring relative light (Lux; 180–1,200 nm) at 60 min intervals during the 4-weeks acclimatization periods, as well as over the course of the 24 h ^14^C incubations (16-17 July and 17-18 September 2020). Average daylight intensity values for June - July and August - September recorded at the top and bottom of the dialysis support frame are shown in Fig. S2.

The ^14^C-radioisotope uptake experiment was carried out on 16 July and 17 September 2020. The ^14^CO_2_ assimilation of every selected microorganism, and a lake water sample collected at the top of the BL, were quantified in sealed glass bottles after a day and night incubation, both in July (at 12.28 m and 12.35 m depth) and September (at 12.63 m and 12.83 m depth).

The dissolved inorganic carbon concentration needed for the calculation^40^ was determined using the CaCO_3_ Merck Spectroquant kit (No. 1.01758.0001) and the Merck Spectroquant Pharo 100 spectrophotometer (Merck, Schaffhausen, Switzerland).

### Light and energy requirements calculations

HOBO light and carbon assimilation data were used to calculate the quantum requirement for the CO_2_ fixation as the ratio between moles of photons absorbed and moles of ^14^CO_2_ fixed by the BL and the dialysis bags pure cultures. Moles were correlated to the surface area (m^2^) of the glass vials used for the ^14^C-incubation to calculate how many moles of photons reached the surface of the vials over the entire light period^41^.

### RNA extraction, sequencing and analysis

Samples for transcriptomic analysis were filtered (Isopore 0.22 mm PC Membrane, diam 25 mm) using a vacuum pump (Vacuumbrand, Wertheim, Germany) connected to a filtration ramp (Pall, Basel, Switzerland) until the filter was completely clogged. Filters were then immediately covered with RNAlater (QIAGEN, Hombrechtikon, Switzerland) for five minutes and stored at -20°C.

RNA was extracted with the RNeasy Plus Universal Mini kit (QIAGEN) following the ‘Purification of total RNA Using the RNeasy Plus Universal mini kit’ protocol for the TissueLyser II, using the complete filter as starting material and a mixture of glass beads of different sizes (0,1 mm, 0,5 mm and 1 mm).

DNase treatment was performed using TURBO DNA-*free* kit (Invitrogen) following the manufacturer’s routine protocol. RNA was quantified fluorometrically at 260/290 nm and 260/230 nm using NanoDrop and the Qubit RNA HS Assay Kit (Invitrogen).

Complementary DNA, PCR and Barcoding of the specimens were performed using the Nanopore SQK-PCB109 kit, according to the accompanying protocol.

For the Oxford Nanopore Technologies (ONT) library preparation, 50-100 fmol in 11 ml of reverse transcribed DNA was used and sequencing performed with an ONT R9.4 flow cell, following the manufacturer’s instructions.

Basecalling and barcoding were performed using the ‘accurate’ option with ONT Guppy software (version 5.0.7), using the command line procedure with the configuration file ‘dna_r9.4.1_450bps_hac.cfg’. RNA quality assessment and postprocessing was performed with ONT Pychopper (v2), followed by running the ONT long-reads pipeline for differential gene expression (DGE) analysis^42^. The bioinformatics workflow employed Snakemake^43^ to run different steps including the following tools: minimap2, salmon, edgeR, DEXSeq and stageR with options (minimap index opts: “”; minimap2 opts: “”; maximum secondary: 100; secondary score ratio: 1.0; salmon libtype: “U”; min samps gene expr: 3; min samps feature expr: 1; min gene expr: 10; min feature expr: 3). NCBI COG (https://www.ncbi.nlm.nih.gov/research/cog-project/) was used to classify the predicted genes into COG-categories.

### Statistical analysis

Data are shown as mean ± SD. All the measurements were taken from distinct samples. Statistical significance was assessed by two-way ANOVA for parametrical data, as indicated; Šidák test was used as a post-hoc test. For multiple comparisons, multiplicity adjusted p-values are indicated in the corresponding figures. Statistical analyses comprising calculation of degrees of freedom were done using GraphPad Prism 9.5.1.

## Results

### Monitoring of the water column

The physico-chemical profile of Lake Cadagno remained substantially unvaried between July and September 2020 (Fig. 1). Parameters measured with the CTD such as dissolved oxygen (DO), light, turbidity, temperature, and conductivity showed little difference between the two periods throughout the water column, except for the location of the turbidity peak and consequently the light intensity reaching the BL. On 16 July 2020, the 1.2 m wide BL lied between 12.18 - 13.36 m depth, with a maximum turbidity value of 17.7 FTU at a depth of about 12.70 m (Fig. 1d), while on 17 September 2020, it was nearly 20 cm wider (1.4 m) and about 70 cm deeper (12.89 – 14.30 m depth) with a maximum turbidity peak of 36.9 FTU almost a meter deeper than in July, at 13.40 m (Fig. 1e). Such difference in depth resulted in a disparate light profile with an intensity reaching the BL top of 0.44 W m^-2^ in September, twice lower than in July with 0.88 W m^-2^ (Fig. 1d, e). Dissolved oxygen showed an irregular profile on 16 July, with the presence of a production peak around 5 m and a minor one around 11 m depth (Fig. 1a), suggesting photosynthetic activity. In September, on the other hand, the profile decreased more linearly without any peak (Fig. 1b). It is relevant to note how in July, and not in September, temperature, conductivity and oxygen at the peak of turbidity showed a homogeneous profile, indicating the presence of a mixed layer due to bioconvection (Fig. 1c). These time points were selected based on the long-term data sets available from the Lake Cadagno monitoring.

**Fig. 1.**
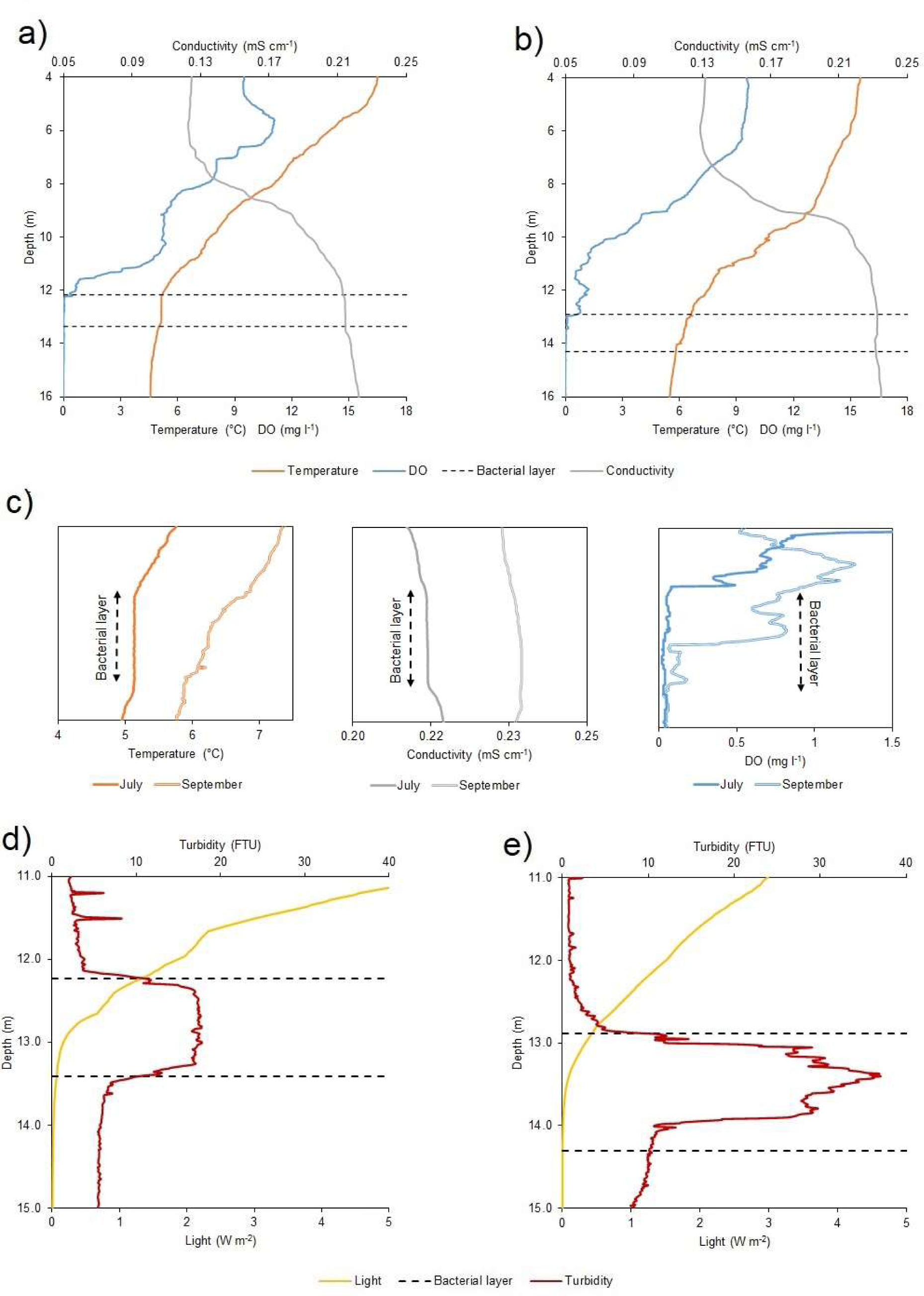
Physicochemical profiles of Lake Cadagno water column. CTD profiles of oxygen (mg l^-1^), temperature (°C), and temperature-corrected (20°C) conductivity (mS cm^-1^) in **a)** July and **b)** September. **c)** Temperature *(left)*, temperature-corrected conductivity *(central)* and DO *(right)* microstructure profiles for the BL and adjacent regions observed on the two different moments of the year 2020. Different light irradiance (W m^-2^) and turbidity (FTU) values within the BL observed between **d)** July and **e)** September. Black dashed lines indicate the BL position on both sampling days (16 July and 17 September 2020).

### Differences in light availability

Data collected with the CTD revealed a difference in the duration and intensity of light in the BL region between July and September. The nearby meteorological station showed an average photoperiod in July of around 16.0 h (05:15 - 21:20) while in September it reduced to about 12.5 h (06:40 – 19:20 h). This resulted in a PAR portion of the average incident radiation below the lake subsurface (1 cm depth) of 100.85 W m^-2^ day^-1^ on 16 July and 43.62 W m^-2^ day^-1^ on 17 September and, consequently, light radiation reaching the top of the BL was higher in July (Tab. 2). Moreover, in September the larger values of observed turbidity (Fig. 1e) determined a greater degree of light attenuation across the BL. As a consequence, the shading effect exerted by the turbidity peak ultimately caused no light to reach the lower part of the BL (Tab. 2).

**Table 1.** Lake Cadagno and BL light regimes (W m^-2^ day^-1^) during the two sampling days.

| Date | Light period (h) | Lake subsurface | Top BL | Bottom BL |
| --- | --- | --- | --- | --- |
| 16.07.20 | 16.0 | 100.85 | 1.44 | 0.06 |
| 17.09.20 | 12.5 | 43.62 | 0.44 | 0.00 |

**Table 2.** July and September ^14^CO_2_ mean assimilation rates (± SD) of the BL microbial community and dialysis bags incubated cultures. Values are reported in ^14^C amol cell^-1^ h^-1^.

|  | July |  | September |  |
| --- | --- | --- | --- | --- |
|  | Day | Night | Day | Night |
| **Bacterial layer** | 6458 ± 159.3 | 1278 ± 62.6 | 1530 ± 43.3 | 1118 ± 27.0 |
| ***C. okenii*** | 5160 ± 481.9 | 2565 ± 140.6 | 981 ± 63.2 | 87 ± 6.0 |
| ***T. syntrophicum*** | 261 ± 4.2 | 104 ± 8.8 | 1144 ± 60.1 | 71 ± 8.0 |
| ***C. phaeobacteroides*** | 8.5 ± 0.7 | 7.8 ± 1.5 | 15 ± 1.0 | 3.0 ± 0.6 |

### Sulfide concentrations across the BL

On 16 July, S^2-^ concentration measured at sunrise, before the light reached the BL, was around 0.14 mg l^-1^ at the top of the BL and 1.88 mg l^-1^ at the bottom, while on 17 September, at the same moment of the day, sulfide concentration at the top was zero (under the detection limit of 0.03 mg l^-1^) and 1.23 mg l^-1^ at the bottom (Fig. 2; yellow bars).

The more homogeneous S^2-^ gradient observed at sunrise in July is also visible when expressing sulfide concentration across the BL as percentage of the concentration at the bottom (Fig. S3).

Interesting to note are the different values of sideward scatter signal (SSC) measured via Flow Cytometry (FCM) (Fig. 2; red and pink bars), which indicates cellular complexity that, for PSB, largely depends on the presence of intracellular sulfur globules^44^. Indeed, in September a decrease in the amount of cell complexity at the top BL was observed in areas gated for *C. okenii* and small-cell PSB in the FCM scatter plots (Fig. S1b). These findings together show that, in proximity of photic zone at the top BL, as sulfide concentration reduces, *C. okenii* and all PSB in general resort to the oxidation of sulfur globules, instead of S^2-^, for their photosynthetic activity.

**Fig. 1.**
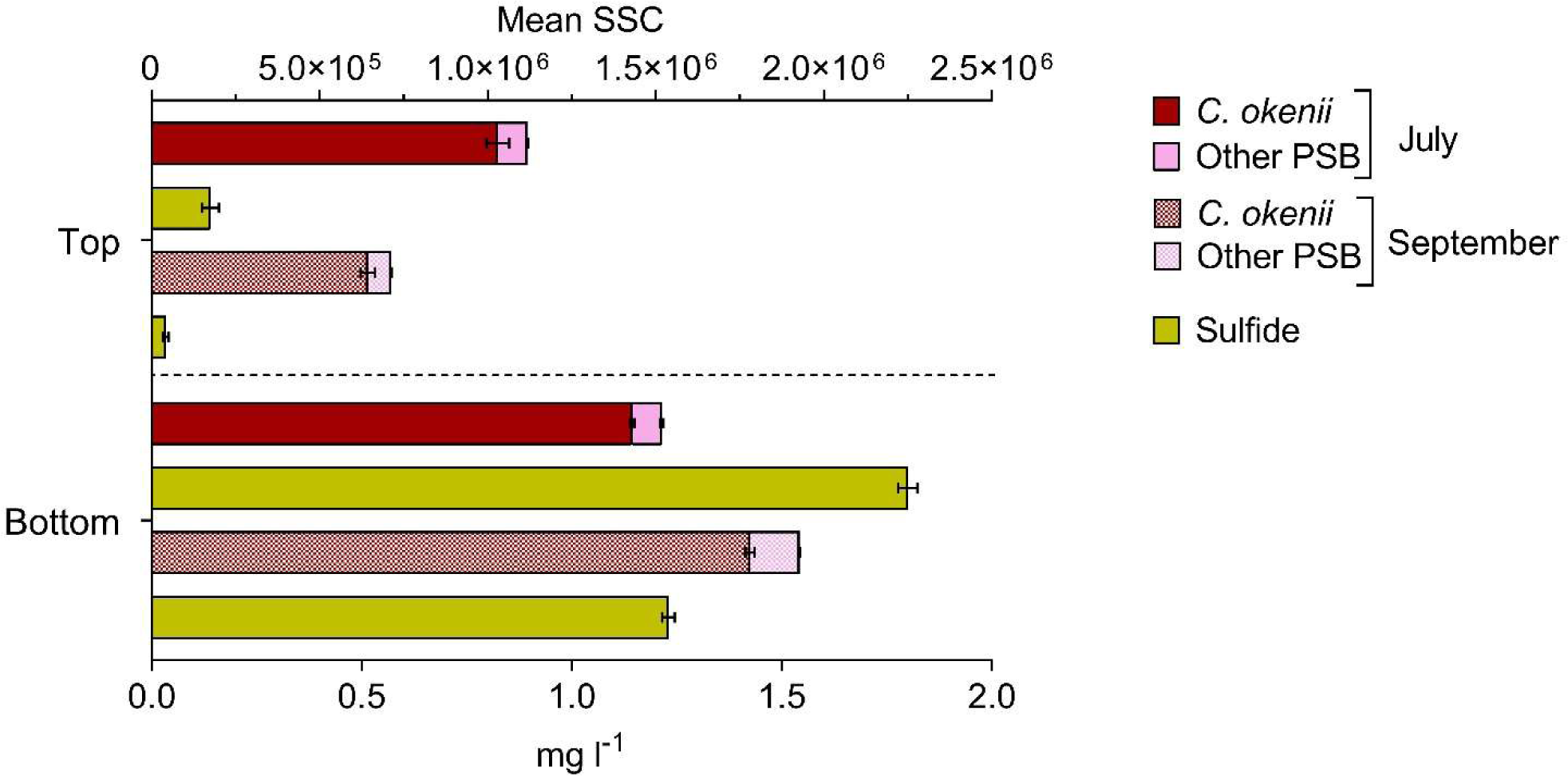
Flow cytometry mean sideward scatter (SSC) of *C. okenii* and small-celled PSB and corresponding sulfide concentrations in the upper and lower part of the bacterial layer on 16 July and 17 September 2020. Error bars represent standard deviation (*n* = 3).

**Fig. 2 a).**
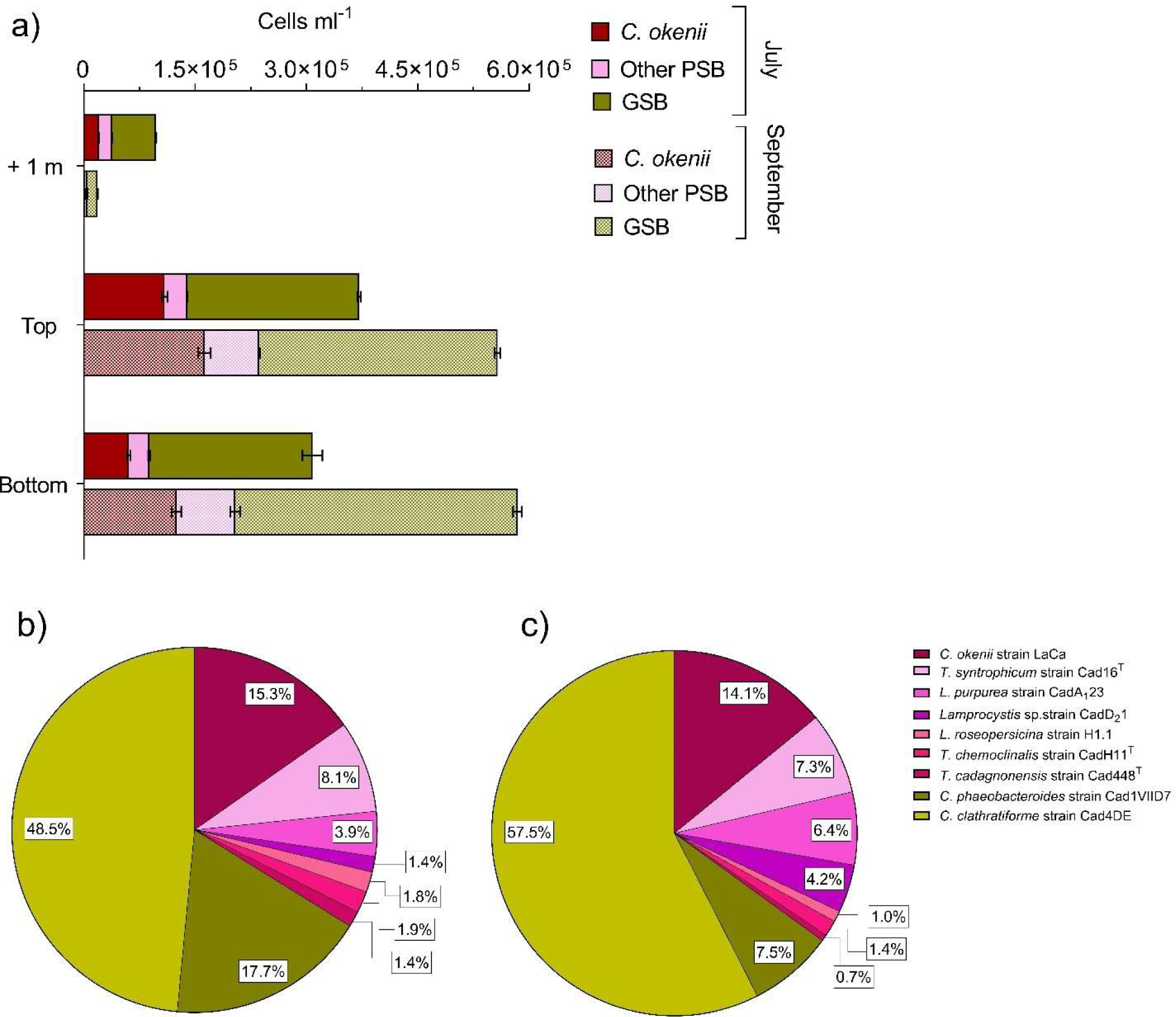
Different abundances of anoxygenic phototrophic sulfur bacteria 1 m above, at the top and bottom of the BL between 16 July 2020 (full color) and 17 September 2020 (speckled color). Cell numbers (x 10^5^ ml^-1^) of all phototrophic sulfur bacteria in the BL obtained by FISH with probes S453F, S453A, S453E, S453H, S453D, S448, CHLP and CHLC at the top of Lake Cadagno BL on **b)** 16 July 2020 and **c)** 17 September 2020. Error bars represent standard deviation (*n* = 3).

### Differences in the BL community

FCM analysis of lake water samples showed high concentrations of photosynthetic microorganisms, within the turbidity peak, in the BL (Fig. 3a).

**Fig. 3.**
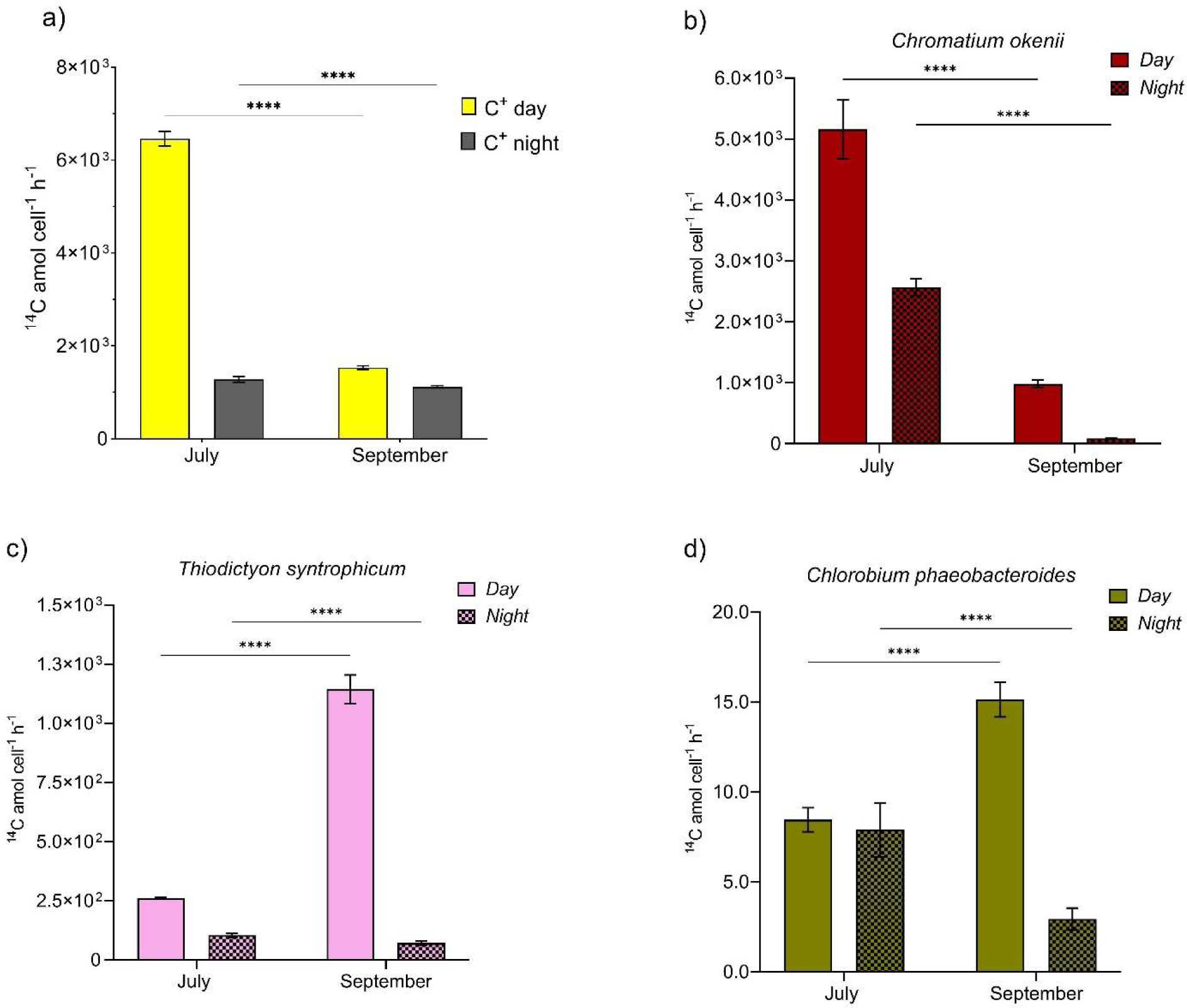
Primary production within the BL. **a**) C-fixation activity of the top BL microbial community in July (left) and September (right), expressed as total ^14^CO_2_ assimilation rate (^14^C amol cell^-1^ h^-1^). **b), c), d)** ^14^CO_2_ uptake by individual pure cultures inside dialysis bags incubated during day (full color bars) and night (dotted color bars) at 12 (July) and 13 (September) m depth. Error bars represent standard deviation (*n* = 3). Error bars represent standard deviation (n = 3). Two-way ANOVA followed by Šidák post hoc comparisons. (****) indicates adjusted *P* values < 0.0001.

In both periods analysed, concentration of large-celled *C. okenii* (approx. length 6 - 8 µm) reached its maximum at the top of the BL, when the turbidity value exceeded 10 FTU, where FCM red fluorescence signal revealed values of 1.09 ± 0.03 x 10^5^ cells ml^-1^ in July and 1.63 ± 0.08 x 10^5^ cells ml^-1^ in September. The relative abundance of *C. okenii* then decreased towards the lower part of the BL with cell concentrations down to 0.61 ± 0.02 x 10^5^ and 1.25 ± 0.06 x 10^5^ cells ml^-1^ in July and September, respectively (Fig. S1b, c; Fig. 3a).

Interestingly, in July during bioconvection, small-celled PSB and GSB were more abundant one meter above the BL, outside the optimal anaerobic zone (5% and 16% of the total counts at the top BL), while in September without bioconvection their numbers above the BL were much lower, accounting for about 1% and 3% of the total cells at the top of the BL, respectively.

Looking in more detail at the species-level composition of the top BL community, large-celled PSB *C. okenii* and all GSB showed a slight decrease in September both about 1.2%, while small-celled PSB had a small increase of 2.4% (Fig. 3). Such increase in PSB abundance, already revealed by FCM counts, was mainly due to PSB *Lamprocystis* sp., strains CadA_1_23 and CadD_2_1 which doubled, from 3.9 to 6.4 %, and quadrupled their counts from 1.4 to 4.2 %, respectively. Among GSB, *Chlorobium clathratiforme* abundance increased by about 9.0%, with a concomitant drop of *C. phaeobacteroides* cell number of 10.2% in September (Fig. 3c).

### Seasonal differences in cell physiology

To investigate the effect of seasonality on the physiology, C-fixation activity, both in presence or absence of light, was measured for the top BL and for single pure cultures of PSB *C. okenii* LaCa and *T. synthophicum*, Cad16^T^ and GSB *C. phaeobacteroides* 1VII D7 incubated in dialysis bags at the corresponding depth of the BL. All samples were added with ^14^CO_2_ and incubated for the whole 16 and 12.5 h light periods, and overnight for the remaining hours, in July and September, respectively, at the corresponding top BL depths (12 and 13 m; Fig. 1d, e). HOBO loggers attached to the structure holding the dialysis bags (top and bottom) confirmed that the ^14^C incubated samples were effectively exposed to the correct light intensity over the course of the experiment (Fig. S2).

On 16 July 2020, the total assimilated ^14^CO_2_ measured from the BL wild population after the diurnal incubation was more than four times higher the daily inorganic carbon fixation rate observed on 17 September 2020 (Fig. 4a; Tab. 2).

Among the dialysis bags cultures incubated during daytime on 16 July, the highest ^14^CO_2_ fixation activity was measured in the large-celled *C. okenii* LaCa (Fig. 4b; Tab. 2). Diurnal assimilation of *T. syntrophicum* Cad16^T^ and *C. phaeobacteroides* 1VII D7 were one and three orders of magnitude lower, respectively (Fig. 4c, d; Tab. 2). In September, a 5-fold decrease in *C. okenii* daytime carbon fixation rate was observed, accompanied by a marked increase in the assimilation activity of both *T. syntrophicum*, which became the major CO_2_ assimilator in September, and *C. phaeobacteroides* (Fig. 4c, d; Tab. 2).

During the night, without any light irradiation, the total inorganic carbon assimilation of the BL decreased slightly from July to September (Fig. 4a; Tab. 2). Night *C. okenii*’s carbon fixation in July was still the highest among the three species, two to three orders of magnitude higher than the values measured for *T. syntrophicum* Cad16^T^ and *C. phaeobacteroides* 1VII D7, respectively (Fig. 4c, d; Tab. 2). Overall, dark CO_2_ assimilation of each single pure culture was higher in July than in September.

Microbial CO_2_ fixation activity in the chemocline of Lake Cadagno was higher in daytime, particularly in July when diurnal carbon assimilation of the BL microbial community was more than double the night one (Fig. 4a). As for pure cultures, it is worth noticing the strong activity of *C. okenii* during both day and night in July, which markedly decreases in September by 81% and 94.3% day and night, respectively.

### Differences in the transcriptome of *C. okenii*

To assess the potential eco-physiological effect of bioconvection on the metabolic activity of its main actor *C. okenii* LaCa, transcriptomic analysis was compared from both dialysis bags cultures of 16 July and 17 September. Transcribed after-night, sunrise-isolated genes from July were compared with those from September, as well as those expressed after-day, sunset, also from July with those from September. After the day, 41 genes resulted upregulated in September, and no one in July, while after night, 45 genes were more expressed in September and 1 in July (Tab. S1, S2).

Of all the 86 genes upregulated from July to September, only in 4 cases it was possible to associate the geneID with a functional category described in COG. The identified categories were (i) signal transduction mechanisms, (ii) cell cycle control, cell division and chromosome partitioning, (iii) translation, ribosomal structure and biogenesis, (iv) lipid transport and metabolism and (v) energy production and conversion.

## Discussion

This study provided for the first time clear indications of positive eco-physiological implications of bioconvection on its promoter, the large-celled PSB *C. okenii*, and the selective metabolic advantage this species gains over other similar microorganisms, namely small-celled PSB and GSB. This was achieved thanks to the possibility of conducting field experiments directly on meromictic Lake Cadagno, where we compared two distinct moments of the year, mid-July, when bioconvection is well present in the BL of the lake, and mid-September, when bioconvection is absent^30, 36^.

To the best of our knowledge, to date we lack compelling evidence of how microbes – in natural settings – leverage bioconvection for eco-physiological benefits. A handful of laboratory-based experiments have suggested that microbes put bioconvection to their benefit for exploiting nutritional microenvironments and improved nutrient absorption efficiency^8, 45, 46^. For example, experimental observations and fluid dynamic simulations have shown that formation of bioconvection can facilitate oxygen transport, which may in turn benefit aerobic microbial communities, as observed in suspensions of *Bacillus subtilis*^47^.

Particularly in natural environments, bioconvection represents an understudied but potentially important mechanism influencing the vertical distribution, and consequently growth and productivity, of the microbial community over different timescales (diurnal and seasonal).

Furthermore, bioconvection could play a role in shaping interspecific interactions and mid- to long-term dynamics in the microbial community in meromictic lakes or, for instance, comprise a metabolic competitive advantage for motile species, overcompensating the substantial energy expenditure resulting from swimming against gravity.

### Chemocline physicochemical parameters

First, we monitored and compared the physico-chemical parameters of Lake Cadagno water column on 16 July and 17 September 2020, which were consistent with past measurements of the same periods^22, 26, 38^. The weather trend for Summer 2020 was within the normal range, with good insolation and little precipitation, with the only exception of a heavy thunderstorm in late August that caused a strong mixing of the mixolimnion, and a consequent increase in light penetration to the BL (Storelli *et al*., manuscript under revision). On 16 July, the water column profile revealed homogeneous temperature and conductivity signatures within the BL, right below the oxic-anoxic interface of the chemocline (Fig. 1c), a proxy of the presence of convective turbulence, which has been shown to be caused by the swimming activity of PSB *C. okenii*^30, 54^. Therefore, the lack of a uniform layer in the CTD profile of 17 September confirmed the absence of bioconvection in late Summer, further highlighting the seasonality of bioconvection, as already observed in previous studies on Lake Cadagno^16, 53^.

Data presented by Sommer *et al*.^30^ showed how, over the course of three summer seasons (August 2014 and 2015, July 2016), bioconvection activity and thickness of the mixed layer in Lake Cadagno positively correlated with *C. okenii* cell concentration. We refer to the authors’ direct numerical simulations (Figure 2 and supporting information Text S3 of the paper) for convincing evidence that the homogeneous layer in Lake Cadagno is due to active biogenic mixing. A detailed explanation of the vertical structure of the mixed layer is provided by Sepúlveda Steiner *et al*.^36^.

Unfortunately, these observations were limited to the time when bioconvection is present. Interestingly, during the period without bioconvection in September, *C. okenii* cell concentration was higher than in July, when bioconvection was well active. This result highlights the complexity of the process driven by the *C. okenii*, seemingly more related to other abiotic or biotic factors investigated in this study, such as light, sulfide or the presence of other phototrophs in the BL, rather than simply to *C. okenii* population size.

### Light period affects bacterial motility

The importance of photoperiod in shaping bacterial motility has been investigated in several studies. For example, experiments on the flagellated microalga *Chlamydomonas reinhardtii*^48^, and other microalgae^49^, revealed that variations in the light-dark period not only markedly affected cell swimming behavior but also influenced cell orientational and gravitactic transport. Similar trends have been presented by other studies on the influence of photoperiod length on the motility rhythm of swimming microorganisms, with important consequences at the population scale^50–52^.

At first the number of cells was identified as a possible factor influencing bioconvection^30^. However, our results suggest photoperiod is a key factor for the onset of bioconvection. In fact, under the shorter light period of September (12.5 h) no mixed layer is observed within the BL, despite the number of *C. okenii* cells was higher than in July (Fig. 1). Further evidence comes from laboratory experiments, where a reduction in the growth rate of *C. okenii* LaCa was observed under a photoperiod of 12/12 h, compared to one of 16/8 h (Di Nezio *et al*., manuscript in preparation). Interestingly, even after changing light intensity from a chemocline-like value of about 4.0 μmol m^-2^ s^-1^ PPFD to a 10-fold increase (about 40 μmol m^-2^ s^-1^ PPFD), the reduction in growth was maintained under the 12/12 h photoperiod, further emphasizing the importance of light period rather than its intensity. Bioconvection in Lake Cadagno has also been reported in the absence of light, suggesting the idea of a continued day-night microbial swimming activity independent of phototaxis^37^. However, it has been suggested that, when nighttime convection occurs, it does so in a fitful fashion and thus only maintaining, rather than expanding, the mixed layer^37^. The ability of *C. okenii* to swim in a coordinated fashion, even without light attraction, therefore suggests the existence of other biophysical mechanisms coordinating movement.

### Bioconvection affects sulfide transport across the bacterial layer

The importance of bioconvection in maintaining chemical gradients across the BL, ensuring the constant influx of key elements, has already been proposed in laboratory studies^35, 53^. Interestingly, the higher S^2-^ concentration across the BL observed in July showed that bioconvection transports and homogenizes sulfide from the depths of the lake (Fig. 2 and S3). In addition to carrying more sulfide, an essential requirement for anoxygenic photosynthesis, bioconvection also promotes the removal of oxygen (Fig. 1c, right), which is chemically reduced by S^2-^. This is further sustained, by higher values of dark CO_2_ fixation as observed in September, without bioconvection, than in July (Fig. 4), when sulfide transport reduced oxygen, reiterating the important role of oxygen in the process of chemosynthesis^28, 29, 38^.

A distinguishing feature of PSB is the production of intracellular sulfur globules (S^0^)^54^, that contribute to determining cell internal complexity, correlating with the SSC parameter measured with FCM^55^. On 16 July at sunrise, when bioconvection was active (Fig. 1c), sulfide was detected up to the oxycline, concurrently with a more homogeneous distribution of intracellular complexity between upper and lower BL (Fig. 2). Conversely, at sunrise on 17 September, with no mixed layer, little S^2-^ was detected at the top BL and it mostly remained confined to the lower part, in concomitance with a more pronounced PSB cell granularity at the bottom BL (Fig. 2). Such S^2-^ distribution was likely caused by microbially-generated convective plumes stirring the water also during night, determining a higher concentration of S^2-^ across the whole BL (Fig. 2). Therefore, the distribution of sulfide throughout the BL determines what mean SSC trend in PSB will exhibit. The higher sulfide concentration in the BL during bioconvection was further pointed out by the oxygen profiles (Fig. 1c, right panel). In fact, coinciding with the turbidity peak (>10 FTU) in July, oxygen was consistently absent along the BL, while in September small amounts of scattered oxygen were measured in the upper BL.

Given these points, it’s clear how bioconvection expand the habitat of the bacterial community, exposing it to light and sweeping S^2-^ along from below.

### Community dynamics in the bacterial layer

The top-bottom small-celled PSB uniform SSC signal in July correlates well with the hypothesis of *C. okenii* dragging along other microorganisms in the BL, namely small-celled PSB and GSB, that are passively moving floating by means of gas vacuoles^21, 24^. In fact, small-celled PSB and GSB populations distribution across the BL highlighted some interesting patterns (Fig. 3a). Interestingly, while *C. okenii* cells were in general more abundant at the top BL on both times of the season, on 16 July, during active bioconvection, we measured a higher number of small-celled PSB and GSB one meter above the BL than on 17 September (without bioconvection). This finding suggested a negative effect toward other microorganisms competing with *C. okenii* for resources such as sulfide and light. Indeed, bioconvection might push many small-celled PSBs and GSBs out of the optimal BL zone in July (+17%), while this is not observed in September (Fig. 3a).

The consequences of bioconvection also appear to affect trophic links, such as grazing by zooplankton. A recent study on the predatory activity of the ciliate *Spirostomum teres,* known to feed on PSB^56^, in Lake Cadagno during the years 2020 and 2021 revealed that, in the presence of bioconvection, the amount of *C. okenii* cells ingested by *S. teres* is significantly lower than when bioconvection is absent (Bolick *et al*., manuscript in preparation).

All together, these observations suggest the effective ecological advantage that *C. okenii* has in producing bioconvection.

### Effect of bioconvection on the primary production

In euxinic environments, light radiation reaching the anaerobic layer is usually positively associated to primary production rates of anaerobic phototrophic sulfur bacteria. The presence of light allows the oxidation of reduced molecules, such as sulfide or sulfur globules, by anaerobic photosynthesis^57^.

Our results indicate that photoperiod plays a relevant role in the ability of *C. okenii* to produce bioconvection. Moreover, a different length of the light period can significantly impact photosynthesis itself, by determining higher or lower light-stimulated rates of inorganic carbon uptake as showed in other photosynthetic bacteria^58–60^, and in PSB and GSB in Lake Cadagno as well. It has been observed that CO_2_ fixation in PSB does not occur at a constant rate throughout the day but reaches the highest values in the first hours of light exposure^61^, compared with the hours of highest light intensity in the afternoon^38^. This trend has also been observed in cyanobacteria^62, 63^. For this reason, adopting whole day and night incubations allowed us to avoid underestimate carbon assimilation activity due to diel cycles.

In this study, the *in situ* daily ^14^C assimilation observed confirmed the strong total inorganic carbon fixation rate in the BL of Lake Cadagno. On 16 July (6.46 ± 0.15 x 10^3^ ^14^C amol cell^-1^ h^-1^), with 16 h of light, diurnal assimilation was more than three times higher than in September, when values reached only 1.53 ± 0.04 x 10^3^ ^14^C amol cell^-1^ h^-1^ after a daylength of 12.5 h (Fig. 4a). This result is certainly strongly influenced by the physiological activity of *C. okenii*. In fact, its intense fixation activity measured in July was much higher than the other microorganisms analyzed (Fig. 4). The situation changed radically in September, when in the absence of bioconvection, *C. okenii* loses dominance over inorganic carbon fixation in favor of the small-celled PSB *T. syntrophicum* Cad16^T^. These findings, in combination with the increase in numbers shown by FISH for all species of small-celled PSB (Fig. 3b), further indicate that bioconvection exerts a negative influence on other microorganisms competing for the same resources as *C. okenii*. Similarly, the higher dark ^14^C assimilation rates observed in July in the BL and in *C. okenii* pure culture (Fig. 4a,b) suggests the persistence of bioconvection during nighttime in Lake Cadagno^37^. In addition, quantum requirements of CO_2_ fixation for each of the three species incubated with carbon-14 were calculated as moles of photons required to assimilate a mole of ^14^CO_2_. The value obtained for *C. okenii* in July (10.6) was similar to that reported by Brune^41^, who calculated quantum requirements of 8.5 - 10.5 quanta per CO_2_ fixed for PSB and 3.3 - 4.5 for GSB, under optimum laboratory conditions, while *T. syntrophicum* and *C. phaeobacteroides* had much higher requirements (23.4 and 74.9, respectively). Under the September light regime, *C. okenii* performed significantly worse (38.5), while *T. syntrophicum* (13.5) and *C. phaeobacteroides* (39.1) had both lower quantum requirements than in July. Applying what observed in the ^14^C-incubated cultures to the broader environmental context of the lake, our data provide enough evidence to sustain that light regime played a key role, as it provided *C. okenii* cells with the energy required for the onset of bioconvection.

### Transcriptomic reveals different gene expression levels

In September the cellular activity of *C. okenii* was much higher compared to July, with the presence of various transcribed genes. The upregulation in *C. okenii* of anoxygenic photosynthesis-related genes in September (Tab. S1) suggests a compensation mechanism for a less efficient photosynthetic activity in the absence of bioconvection and under less favorable light conditions ^64, 65^. Data from transcriptomic analyses on *C. okenii* corroborated these findings showing higher expression levels of genes involved in the photoautotrophic sulfur oxidation when in absence of bioconvection.

However, despite the significant contributions transcriptomics has made to the field of microbial ecology, some limitations have emerged in the application of this techniques to the study of physiological responses to the environment^66^. The main constraints are related to the fact that genes with relevant impact on fitness are rare and therefore difficult to detect by transcript analysis, and besides, the relationship between gene expression and fitness is often dubious. Another limit is that fitness is mostly determined by protein activity and the amount of mRNA is a poor indicator of the amount of protein^67^. Nonetheless, (meta)transcriptomics has the potential of providing valuable insights into environmental microbial communities.

### Conclusions

In this paper, we combined biological, chemical and physical factors to elucidate how bioconvection shape the main eco-physiological traits for the phototrophic sulfur bacteria community inhabiting the BL of Lake Cadagno. We first report the key role of the photoperiod length by comparing two different period of measurements. Moreover, we showed how the presence of bioconvection contributes to maintaining a sulfide gradient across the BL, thereby promoting oxygen removal and avoiding consumption of intracellular storage substances such as sulfur globules, requiring a greater energy investment (also shown by the high transcriptional activity in the absence of bioconvection). It is also interesting to note that bioconvection negatively affects the fitness of the other ecological competitors of *C. okenii*, namely small-celled PSB and GSB, present in the BL. Overall, our combined data suggest that *C. okenii* is able to gain a competitive advantage over other non-motile phototrophic sulfur bacteria in the quest for the optimal environmental conditions by producing mixed layers through bioconvection.

Nevertheless, despite this study provides evidence of the eco-physiological effects of bioconvection in an environmental setting, its consequences on the microenvironmental conditions and the other (micro)organisms involved need to be substantiated with further studies. In particular, the role of bioconvection on the transport of sulfide across the BL, and further insights on its production by sulfate-reducing bacteria in the monimolimnion, require a more detailed investigation. Impacts of bioconvection might also extend outside the mixed layer, influencing the interaction between phototrophic sulfur bacteria and the zooplankton living just above the bacterial layer, e.g., in terms of predation and/or distribution patterns. Lastly, a deeper understanding of the motility mechanisms of *C. okenii* triggering bioconvection at the single-cell level will help unravelling the nature of this multi-scale process.

## Acknowledgments

We thank G. Ranieri and M. Thelen for technical assistance with ^14^C experiment analyses; M. Tonolla and F. Danza for fruitful discussion and constructive comments on the manuscript; the Alpine Biology Centre Foundation (CBA) for laboratory facilities and housing. Funding was provided by the Swiss National Science Foundation (grant number 315230–179264) and by the Institute of Microbiology (IM) of the University of Applied Sciences and Arts of Southern Switzerland (SUPSI). A. Sengupta thanks the ATTRACT Investigator Grant (No. A17/MS/11572821/MBRACE) and FNR-CORE Grant (No. C19/MS/13719464/TOPOFLUME/Sengupta) from the Luxembourg National Research Fund for supporting this work.

## Author contributions

FDN, SR, ABD, and NS designed research; FDN, SR, ABD, and NS collected samples; FDN and SR performed laboratory work; FDN, SR, OSS and ABD analyzed data; FDN and NS wrote the paper. All authors developed the concepts and hypotheses covered in this work, and have contributed to the revisions of the manuscript.

