## Supplementary material for "Motile bacteria leverage bioconvection for eco-physiological benefits in a natural aquatic environment"

[1]University of Applied Sciences and Arts of Southern Switzerland (SUPSI), Department of Environment, Constructions and Design, Institute of Microbiology, Via Flora Ruchat-Roncati 15, 6850 Mendrisio, Switzerland. [2]University of Geneva, Department of Plant Sciences, Boulevard d'Yvoy 4, 1205 Geneva, Switzerland. [3]Alpine Biology Center Foundation, Via Mirasole 22A, 6500 Bellinzona, Switzerland. [4]Swiss Federal Institute of Aquatic Science and Technology (Eawag), Department of Surface Waters - Research and Management, Seestrasse 79, 6047 Kastanienbaum, Switzerland. [5]Civil & Environmental Engineering, University of California – Davis, 3155 Ghausi Hall, Davis, CA 95616, USA. [6]Physics of Living Matter, Department of Physics and Materials Science, 162A Avenue de la Faïencerie, L-1511 Luxembourg City, Luxembourg. *

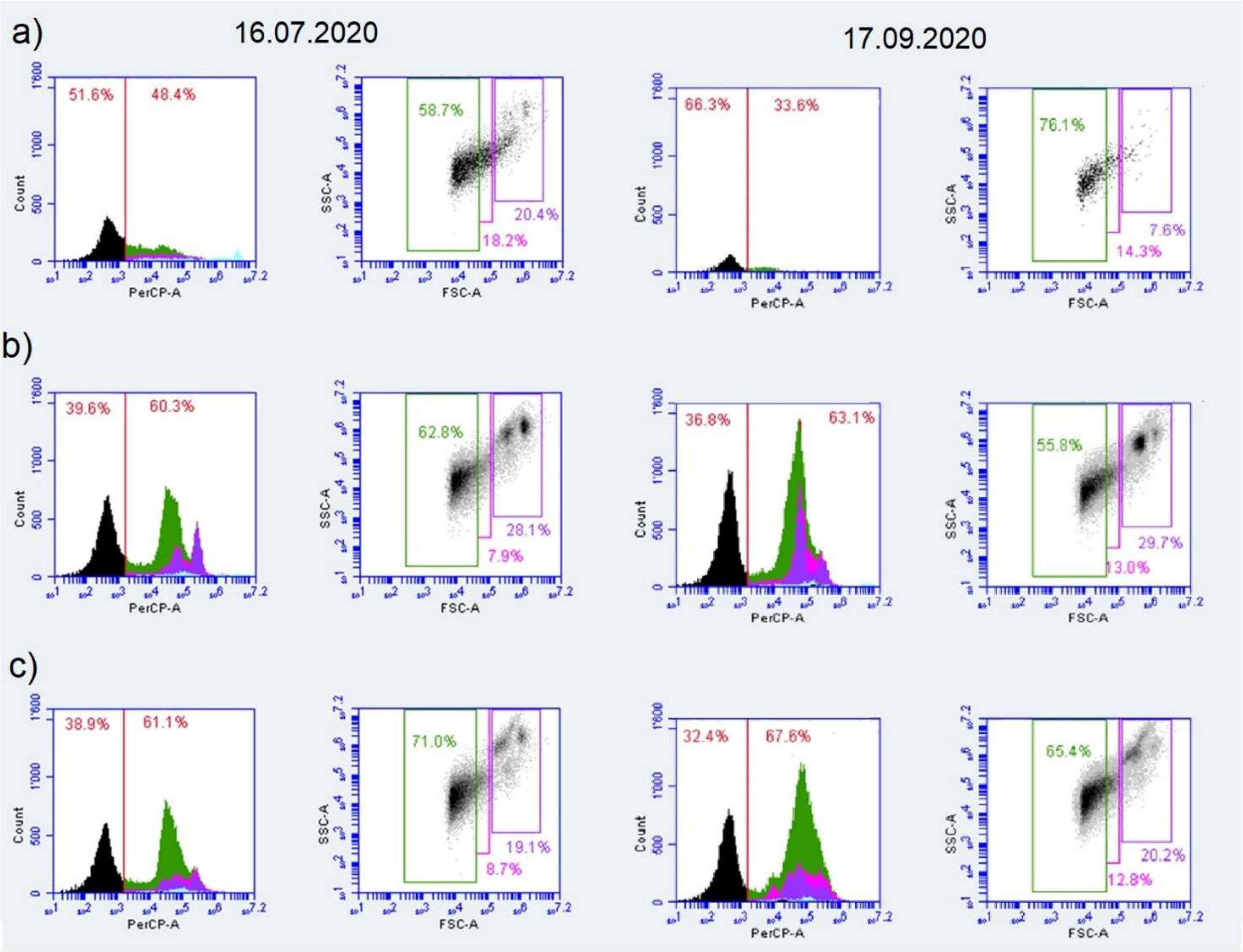


**Fig. S1** Flow-cytometry identification and relative abundances of phototrophic populations at **a)** 1 m above, **b)** top and **c)** bottom bacterial layer on 16 July and 17 September 2020. For each date: (*left*) histograms count *vs* red fluorescence (logarithmic), red line (PerCP-A > 1 100) separates chlorophyll-pigmented cells from non-autofluorescent cells with relative percentages of the total counts; (*right*) scatter plots SSC *vs* FSC with relative percentages of the total chlorophyll-pigmented cells. Purple color identifies PSB *C. okenii* population (8 - 10 μm), pink defines small-celled PSB (2 - 4 μm) and green refers to GSB cells (0.1 - 1 μm),

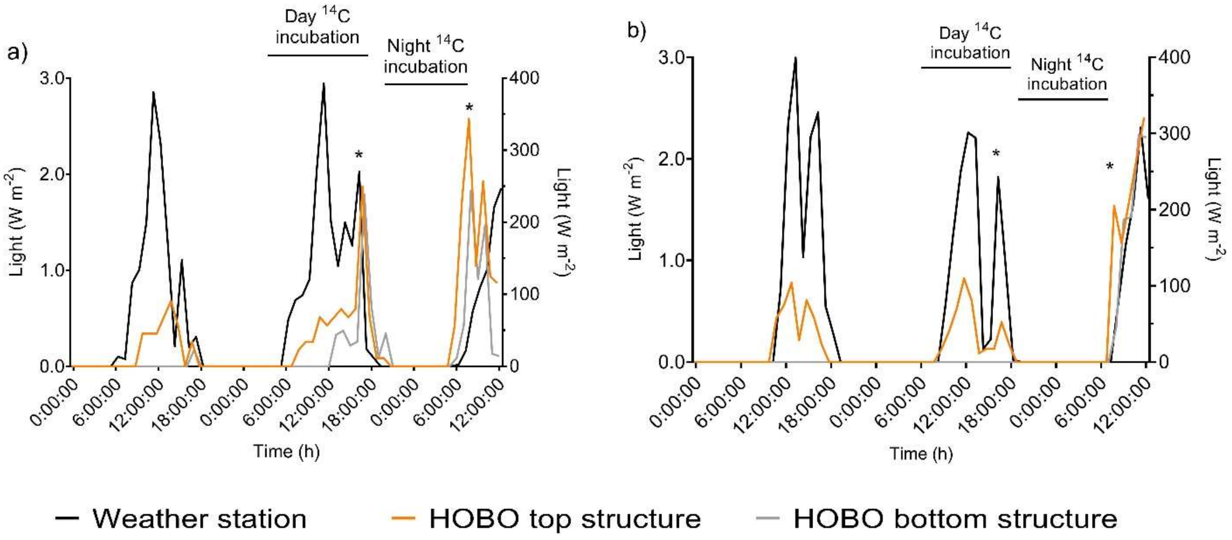


**Fig. S2** Net radiation intensity recorded by the weather station and HOBO loggers attached at the top and bottom of the dialysis bags structure during **a)** 15-17 July 2020 and **b)** 16-18 September 2020. Asterisks indicate the moment the structure was retrieved at the end of the $^{14}$C incubation periods (corresponding HOBO light values divided by 100).

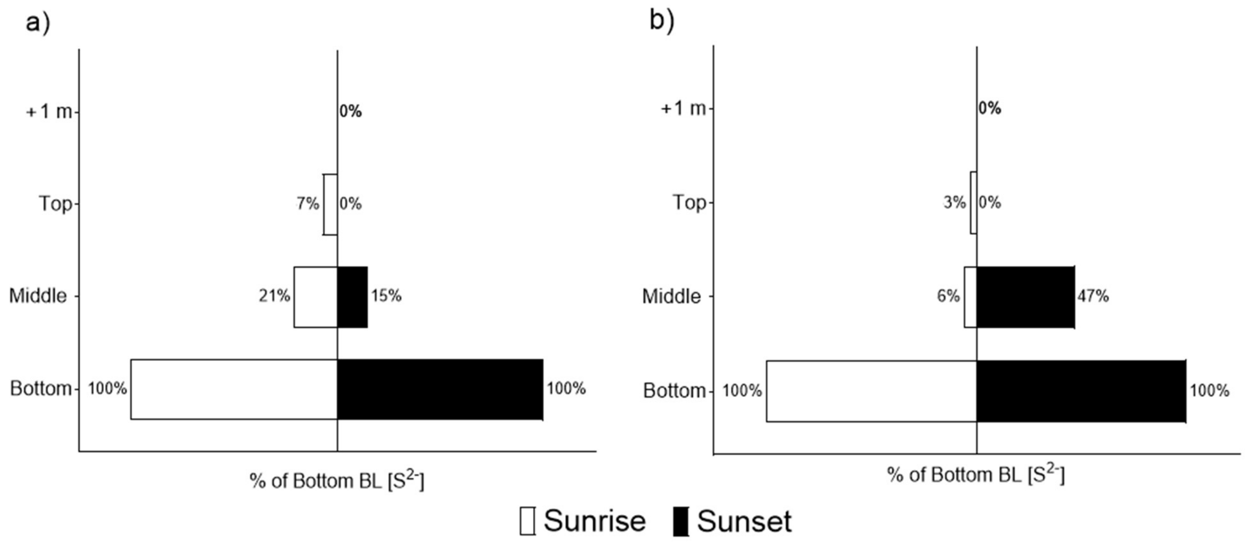


**Fig. S3** Sulfide concentration across the BL expressed as percentage of the concentration at the bottom, at sunrise and sunset on **a)** 16 July 2020 and **b)** 17 September 2020.

**Table S1** Oligonucleotide probes for the in situ hybridization assay (FISH).

| Probe | Target | Sequence | Formamide % |
|---|---|---|---|
| Cmok453[40] | *Chromatium okenii* (DSM 169) 16S rRNA, pos.453-479 | AGCCGATGGGTATTAACCACCAGGTT | 30 |
| S453F[40] | *Thiodyction synthrophicum* 16S rRNA, pos. 453-479 | CGGCGTCGCTGCGTCAGG | 40 |
| S453A[40] | *Lamprocystis purpurea* (DSM 4197) 16S rRNA, pos. 453-479 | TCGCCCAGGGTATTATCCCAAACGAC | 40 |
| S453E[40] | *Lamprocystis roseopersicina* (DSM 229) 16S rRNA, pos. 453-479 | CATTCCAGGGTATTAACCCAAAATGC | 30 |
| S453H[19] | *Thiocystis chemoclinalis* 16S rRNA, pos. 453-478 | GACGGAACGGTATTAACGCCCCGCTT | 10 |
| S448[19] | *Thiocystis cadagnonensis* 16S rRNA, pos. 448-468 | CGGCGTCGCTGCGTCAGG | 25 |
| S453D[40] | Clone 261 from Lake Cadagno 16S rRNA, pos. 453-479 | CAGCCCAGGGTATTAACCCAAGCCGC | 40 |
| CHLP[41] | *Chlorobium phaeobacteroides* (DSM 266), 16S rRNA, pos. 441-464 | AAATCGGGATATTCTTCCTCCAC | 20 |
| CHLC[19] | *Chlorobium clathratiforme* (DSM 5477), 16S rRNA, pos. 190-211 | GGCAGAACAACCATGCGATTGT | 20 |
| DSC441 + DSC213[42] | *Desulfocapsa thiozymogenes* 213-230 441-459 | ATTACACTTCTTCCCATCC (DSC441) CCTCCCTGTACGATAGCT (DSC213) | |

**Table S2** Day upregulated genes of *C. okenii* in September.

| Gene ID | Gene symbol | Description | p value | $Log_2FC$ |
|---|---|---|---|---|
| lcl\|NZ_PPGH01000038.1_cds_WP_105074862.1_3481 | | DUF4342 domain-containing protein | 8.35E-06 | 10.137 |
| lcl\|NZ_PPGH01000034.1_cds_WP_105073399.1_1552 | | light-harvesting protein | 2.59E-06 | 9.872 |
| lcl\|NZ_PPGH01000018.1_cds_958 | | transposase | 2.78E-06 | 8.000 |
| lcl\|NZ_PPGH01000037.1_cds_WP_105074161.1_2503 | *acp* | acyl carrier protein | 6.71E-06 | 7.948 |
| lcl\|NZ_PPGH01000018.1_cds_WP_172452521.1_751 | *csd* | cold shock domain-containing protein | 2.28E-05 | 7.824 |
| lcl\|NZ_PPGH01000010.1_cds_351 | | cyclic nucleotide-binding domain-containing protein | 3.11E-05 | 7.474 |
| lcl\|NZ_PPGH01000038.1_cds_3472 | *aprA* | adenylyl-sulfate reductase subunit alpha | 5.48E-05 | 7.177 |
| lcl\|NZ_PPGH01000037.1_cds_WP_105074694.1_3335 | *prfB* | peptide chain release factor 2 | 7.00E-05 | 7.047 |
| lcl\|NZ_PPGH01000037.1_cds_2771 | | signal recognition particle protein | 9.15E-05 | 6.958 |
| lcl\|NZ_PPGH01000037.1_cds_3027 | *alats* | alanine--tRNA ligase | 9.30E-05 | 6.781 |
| lcl\|NZ_PPGH01000037.1_cds_WP_105074364.1_2797 | *hns* | H-NS histone family protein | 9.52E-05 | 6.720 |
| lcl\|NZ_PPGH01000038.1_cds_3625 | *infA1* | translation initiation factor IF-1 | 1.14E-04 | 6.701 |
| lcl\|NZ_PPGH01000034.1_cds_WP_105073506.1_1611 | *sgp* | sulfur globule protein CV1 | 1.36E-04 | 6.643 |
| lcl\|NZ_PPGH01000037.1_cds_WP_105074683.1_3317 | *ompH* | OmpH family outer membrane protein | 1.37E-04 | 6.623 |
| lcl\|NZ_PPGH01000037.1_cds_WP_146108813.1_2502 | *fabG* | 3-oxoacyl-ACP reductase FabG | 1.53E-04 | 6.613 |
| lcl\|NZ_PPGH01000035.1_cds_WP_105073908.1_2221 | *Scp2d1* | SCP2 sterol-binding domain-containing protein | 1.56E-04 | 6.579 |
| lcl\|NZ_PPGH01000034.1_cds_WP_105073369.1_1505 | *rpoN* | RNA polymerase factor sigma-54 | 1.75E-04 | 6.553 |
| lcl\|NZ_PPGH01000035.1_cds_WP_105073703.1_1935 | *rbcs* | ribulose-bisphosphate carboxylase | 3.09E-04 | 6.516 |
| lcl\|NZ_PPGH01000034.1_cds_WP_105073474.1_1659 | | ATPase | 3.29E-04 | 6.516 |
| lcl\|NZ_PPGH01000037.1_cds_3107 | *holA* | DNA polymerase III subunit delta | 3.48E-04 | 6.484 |
| lcl\|NZ_PPGH01000024.1_cds_WP_105073055.1_62 | *ProQ/FinO* | ProQ/FinO family protein | 3.63E-04 | 6.475 |
| lcl\|NZ_PPGH01000018.1_cds_WP_105072968.1_971 | *ompA* | OmpA family protein | 5.80E-04 | 6.442 |
| lcl\|NZ_PPGH01000038.1_cds_WP_105074959.1_3628 | | carboxynorspermidine decarboxylase | 5.74E-04 | 6.434 |
| lcl\|NZ_PPGH01000035.1_cds_WP_105073619.1_1829 | *rps5* | 30S ribosomal protein S5 | 5.84E-04 | 6.286 |
| lcl\|NZ_PPGH01000018.1_cds_863 | | bifunctional D-glycero-beta-D-manno-heptose-7-phosphate kinase/D-glycero-beta-D-manno-heptose 1-phosphate adenylyltransferase HldE | 6.21E-04 | 6.254 |
| lcl\|NZ_PPGH01000037.1_cds_2566 | *tssC* | type VI secretion system contractile sheath large subunit | 6.05E-04 | 6.247 |
| lcl\|NZ_PPGH01000037.1_cds_WP_105074184.1_2529 | *nifK* | nitrogenase molybdenum-iron protein subunit beta | 6.54E-04 | 6.224 |
| lcl\|NZ_PPGH01000035.1_cds_WP_105073969.1_2313 | | 50S ribosome-binding GTPase | 7.09E-04 | 6.222 |
| lcl\|NZ_PPGH01000035.1_cds_WP_105073877.1_2171 | *rmlB* | 23S rRNA (guanosine (2251) -2'-O) -methyltransferase RlmB | 7.08E-04 | 6.195 |
| lcl\|NZ_PPGH01000035.1_cds_WP_105073901.1_2213 | *infA2* | translation initiation factor IF-2 | 7.20E-04 | 6.095 |
| lcl\|NZ_PPGH01000035.1_cds_WP_105073876.1_2170 | *rnr* | ribonuclease R | 4.30E-05 | 6.091 |

| | | | | |
|---|---|---|---|---|
| lcl\|NZ_PPGH01000034.1_cds_1423 | *fbp* | class 1 fructose-bisphosphatase | 3.87E-05 | 6.089 |
| lcl\|NZ_PPGH01000034.1_cds_WP_105073368.1_1504 | *raiA* | ribosome-associated translation inhibitor RaiA | 7.26E-05 | 6.076 |
| lcl\|NZ_PPGH01000037.1_cds_2721 | *cheB_2* | chemotaxis response regulator protein-glutamate methylesterase | 1.62E-04 | 6.074 |
| lcl\|NZ_PPGH01000037.1_cds_WP_105074636.1_3237 | *pstS_3* | PstS family phosphate ABC transporter substrate-binding protein | 1.68E-04 | 6.049 |
| lcl\|NZ_PPGH01000037.1_cds_3212 | *tusA* | sulfurtransferase TusA family protein | 1.95E-04 | 6.033 |
| lcl\|NZ_PPGH01000035.1_cds_WP_105074045.1_2164 | *infA3* | translation initiation factor IF-3 | 2.53E-04 | 6.028 |
| lcl\|NZ_PPGH01000035.1_cds_2175 | | polyprenyl synthetase family protein | 9.09E-04 | 6.026 |
| lcl\|NZ_PPGH01000037.1_cds_WP_105074266.1_2651 | | DUF4347 domain-containing protein | 6.08E-04 | 5.101 |
| lcl\|NZ_PPGH01000035.1_cds_WP_105073750.1_2000 | *tssH* | type VI secretion system ATPase TssH | 8.71E-04 | 4.843 |

**Table S3** Night upregulated genes of *C. okenii* in September.

| GeneID | Gene symbol | Description | p value | $Log_2FC$ |
|---|---|---|---|---|
| lcl\|NZ_PPGH01000010.1_cds_WP_105072512.1_328 | *rps16* | 30S ribosomal protein S16 | 8.34E-10 | 7.669 |
| lcl\|NZ_PPGH01000037.1_cds_3212 | *tusA* | sulfurtransferase TusA family protein | 7.12E-07 | 6.806 |
| lcl\|NZ_PPGH01000035.1_cds_WP_105073947.1_2285 | | PEP-CTERM sorting domain-containing protein | 7.43E-07 | 6.805 |
| lcl\|NZ_PPGH01000013.1_cds_WP_105072622.1_579 | | DUF2760 domain-containing protein | 2.51E-06 | 6.625 |
| lcl\|NZ_PPGH01000037.1_cds_2771 | *srp54* | signal recognition particle protein | 9.33E-06 | 6.419 |
| lcl\|NZ_PPGH01000034.1_cds_WP_105073398.1_1551 | | light-harvesting protein | 9.34E-06 | 6.413 |
| lcl\|NZ_PPGH01000016.1_cds_714 | *ATPsynB* | V-type ATP synthase subunit B | 1.14E-05 | 6.393 |
| lcl\|NZ_PPGH01000034.1_cds_WP_105073483.1_1332 | *rsmA* | 16S rRNA (adenine(1518)-N(6)/adenine(1519)-N(6))-dimethyltransferase RsmA | 1.99E-05 | 6.310 |
| lcl\|NZ_PPGH01000034.1_cds_1344 | *surA* | SurA N-terminal domain-containing protein | 1.91E-05 | 6.299 |
| lcl\|NZ_PPGH01000037.1_cds_WP_105074511.1_3024 | *csrA* | carbon storage regulator CsrA | 5.52E-05 | 6.117 |
| lcl\|NZ_PPGH01000037.1_cds_WP_105074528.1_3057 | *ftsZ* | cell division protein FtsZ | 5.98E-05 | 6.112 |
| lcl\|NZ_PPGH01000035.1_cds_WP_105073969.1_2313 | | 50S ribosome-binding GTPase | 6.24E-05 | 6.106 |
| lcl\|NZ_PPGH01000035.1_cds_WP_105073615.1_1824 | *rps13* | 30S ribosomal protein S13 | 6.53E-05 | 6.087 |
| lcl\|NZ_PPGH01000035.1_cds_2231 | *rsxC* | electron transport complex subunit RsxC | 7.56E-05 | 6.054 |
| lcl\|NZ_PPGH01000010.1_cds_351 | | cyclic nucleotide-binding domain-containing protein | 8.38E-05 | 6.049 |
| lcl\|NZ_PPGH01000035.1_cds_WP_105073637.1_1850 | *rps7* | 30S ribosomal protein S7 | 9.86E-05 | 6.019 |
| lcl\|NZ_PPGH01000037.1_cds_WP_105072420.1_2775 | *shmt1* | serine hydroxymethyltransferase | 1.28E-04 | 5.975 |
| lcl\|NZ_PPGH01000035.1_cds_WP_105073866.1_2159 | *ihfA* | integration host factor subunit alpha | 1.51E-04 | 5.929 |
| lcl\|NZ_PPGH01000035.1_cds_WP_105074045.1_2164 | *infA3* | translation initiation factor IF-3 | 1.04E-11 | 5.922 |
| lcl\|NZ_PPGH01000038.1_cds_3625 | *infA1* | translation initiation factor IF-1 | 1.52E-11 | 5.878 |
| lcl\|NZ_PPGH01000038.1_cds_3472 | *aprA* | adenylyl-sulfate reductase subunit alpha | 2.22E-04 | 5.858 |
| lcl\|NZ_PPGH01000020.1_cds_1066 | | NADP-dependent isocitrate dehydrogenase | 5.79E-04 | 5.649 |
| lcl\|NZ_PPGH01000035.1_cds_WP_105073867.1_2160 | *pheT* | phenylalanine--tRNA ligase subunit beta | 5.67E-04 | 5.645 |
| lcl\|NZ_PPGH01000034.1_cds_WP_105073477.1_1663 | | glucokinase | 6.73E-04 | 5.608 |
| lcl\|NZ_PPGH01000039.1_cds_WP_105074862.1_3643 | | DUF4342 domain-containing protein | 3.49E-10 | 5.532 |
| lcl\|NZ_PPGH01000037.1_cds_WP_105074512.1_3025 | | aspartate kinase | 9.37E-04 | 5.522 |
| lcl\|NZ_PPGH01000018.1_cds_WP_105072748.1_758 | *ilvB1* | biosynthetic-type acetolactate synthase large subunit | 5.91E-12 | 5.357 |
| lcl\|NZ_PPGH01000037.1_cds_WP_105074694.1_3335 | *prfB* | peptide chain release factor 2 | 3.54E-08 | 5.002 |
| lcl\|NZ_PPGH01000034.1_cds_WP_105073474.1_1659 | | ATPase | 5.91E-06 | 4.336 |
| lcl\|NZ_PPGH01000013.1_cds_WP_105072630.1_587 | *nuoD* | NADH-quinone oxidoreductase subunit D | 3.43E-05 | 4.060 |

| | | | | |
|---|---|---|---|---|
| lcl\|NZ_PPGH01000035.1_cds_WP_105073876.1_2170 | | ribonuclease R | 3.03E-10 | 4.007 |
| lcl\|NZ_PPGH01000037.1_cds_WP_105074161.1_2503 | | acyl carrier protein | 1.08E-06 | 3.636 |
| lcl\|NZ_PPGH01000037.1_cds_WP_105074110.1_2435 | | 4-(cytidine 5'-diphospho)-2-C-methyl-D-erythritol kinase | 4.51E-05 | 3.445 |
| lcl\|NZ_PPGH01000034.1_cds_1423 | *fbp* | class 1 fructose-bisphosphatase | 5.22E-05 | 3.413 |
| lcl\|NZ_PPGH01000037.1_cds_WP_105074768.1_2799 | | RNA-binding protein | 2.01E-04 | 3.203 |
| lcl\|NZ_PPGH01000037.1_cds_WP_105074636.1_3237 | *pstS* | PstS family phosphate ABC transporter substrate-binding protein | 7.81E-07 | 3.199 |
| lcl\|NZ_PPGH01000035.1_cds_WP_105073752.1_2003 | *rps1* | 30S ribosomal protein S1 | 7.88E-06 | 3.165 |
| lcl\|NZ_PPGH01000037.1_cds_WP_105074253.1_2634 | | chaperonin GroEL | 9.54E-05 | 3.022 |
| lcl\|NZ_PPGH01000035.1_cds_WP_105073619.1_1829 | *rps5* | 30S ribosomal protein S5 | 9.34E-04 | 2.932 |
| lcl\|NZ_PPGH01000021.1_cds_WP_105072992.1_1078 | *ProQ/FinO* | ProQ/FinO family protein | 7.55E-07 | 2.901 |
| lcl\|NZ_PPGH01000039.1_cds_3634 | | adenylyl-sulfate reductase subunit beta | 5.52E-05 | 2.896 |
| lcl\|NZ_PPGH01000035.1_cds_2175 | | polyprenyl synthetase family protein | 6.89E-04 | 2.683 |
| lcl\|NZ_PPGH01000018.1_cds_WP_172452521.1_751 | *csd* | cold shock domain-containing protein | 5.27E-04 | 2.163 |
| lcl\|NZ_PPGH01000035.1_cds_WP_197711031.1_1874 | | transposase | 3.41E-04 | -3.063 |
